# Comprehensive assembly of monoclonal and mixed antibody sequences

**DOI:** 10.1101/2024.08.09.607415

**Authors:** Wenbin Jiang, Yueting Xiong, Jin Xiao, Jingyi Wang, Zhenjian Jiang, Ling Luo, Quan Yuan, Ningshao Xia, Rongshan Yu

**Affiliations:** National Institute for Data Science in Health and Medicine, State Key Laboratory of Vaccines for Infectious Diseases, Xiang An Biomedicine Laboratory, School of Public Health, Xiamen University, Xiamen, Fujian, China; School of Informatics, Xiamen University, Xiamen, Fujian, China; Aginome Scientific, Xiamen, Fujian, China; Department of Pathology, Zhongshan Hospital, Fudan University (Xiamen Branch), Xiamen, Fujian, China

**Keywords:** Antibody, *De novo* sequencing, Assembly, High throughput

## Abstract

The elucidation of antibody sequence information is crucial for understanding antigen binding and advancing therapeutic and research applications. However, complete *de novo* assembly of monoclonal antibody sequences remains challenging due to accuracy and robustness limitations. To address this issue, we introduce Fusion, an innovative *de novo* assembler that integrates overlapping peptides and template information into complete sequences using a beam search strategy. We demonstrate Fusion’s performance by reconstructing multiple human and murine antibodies with highest accuracy (100% and over 99%, respectively). Biological validation of the recombinantly expressed AFS98 antibody with unknown sequences further supports its effectiveness. Furthermore, current methods are applicable only to traditional monoclonal antibody sequencing assembly, presenting a significant bottleneck in achieving higher throughput. In contrast, Fusion can assemble peptide sequences from mixtures of two or three monoclonal antibodies into complete individual sequences with the same accuracy as traditional sequencing, significantly enhancing throughput. To our knowledge, this is the first study enabling high-throughput sequencing of multiple antibodies using only bottom-up mass spectrometry. The duration, expense, and reagent consumption of mass spectrometry detection are comparable to those required for sequencing a single monoclonal antibody. In summary, Fusion’s superior performance in handling the complex antibody sequencing represents a significant advancement in antibody research.

## 1 Introduction

Antibodies, in the form of large Y-shaped immunoglobulins, play a crucial role in the immune system by identifying and neutralizing foreign entities such as pathogenic bacteria and viruses [1]. In addition to directly neutralizing pathogens, antibodies coordinate a broader immune response by recruiting effector functions for mechanisms, such as antibody-dependent cellular cytotoxicity and complement-dependent cytotoxicity [2]. Furthermore, antibodies produced in response to an infection can remain in circulation for prolonged durations, allowing for rapid and strong responses upon reencountering the same pathogen due to the memory B-cells’ ability to recall [3]. The multifaceted functionality of antibodies underscores their critical importance across a broad spectrum of disciplines, encompassing immunology, clinical chemistry, biochemistry, therapeutics, and medicine [4–6]. As a result, they have become invaluable tools in numerous research and practical applications.

In humans and most mammals, an antibody molecule is composed of two identical heavy chains and two identical light chains. These chains are linked by disulfide bonds, with each chain consisting of a variable region for antigen binding and a constant region contributing to oligomerization, complement activation, and receptor binding on immune cells [7]. Each chain is encoded by up to four separate gene segments through somatic recombination of the Variable, Diversity (only in heavy chain), Joining, and Constant segments. The vast sequence diversity in antibodies necessary for binding a wide range of antigens arises from V-(D)-J-C recombination, hypermutation, and pairings between heavy and light chains. The specificity (recognition) and affinity (binding) of the antibodies to their antigens are primarily governed by three complementarity determining regions (CDRs). Specifically, CDR1 and CDR2 are encoded by the V segment, while CDR3, which exhibits the greatest variability, is encoded in the V-(D)-J junction.

The retrieval of sequence information for antibodies is essential for understanding the structural underpinnings of antibody-antigen binding, recognition, and interaction [8]. With the advancement of next-generation protein sequencing technologies, tandem mass spectrometry (MS/MS) has emerged as a powerful technique for determining the amino acid sequence of an antibody [9–11]. In standard bottom-up mass spectrometry, antibody samples undergo multiple enzymatic digestions into shorter peptides to facilitate MS/MS analysis [12]. Subsequently, the obtained spectra require computational analysis and statistical extrapolation. For novel or unknown antibodies, *de novo* peptide sequencing is commonly and initially used to obtain sequence information [13]. This method directly identifies fragmented peptides by computing mass differences of ions from related MS/MS spectra, without relying on any sequence database. Notably, *de novo* peptide sequencing has improved in recent years with multiple published methods showing promising results for predicting peptide sequences [14–17].

However, the complete assembly of antibody sequences presents another challenging task. Current methods for this are limited and can be categorized into two types based on their use of database templates. The ALPS algorithm [18], a greedy algorithm-based *de novo* assembly system, utilizes *de novo* peptides, sequence confidence scores, and database homology search information to build a weighted de Bruijn graph for assembling monoclonal antibody sequences. This approach effectively assembles local fragments of the antibody sequence with high accuracy, owing to the consensus sequence derived from the overlap of k-mers. But this template-independent technique is unable to fully assemble a whole chain due to inherent errors in *de novo* peptide sequencing. Additionally, ALPS struggles to differentiate between light and heavy chains in mixed data. On the other hand, the Stitch algorithm [7] maps proteomic short reads to available germline templates for fully reconstructing monoclonal antibody sequences. Nevertheless, Stitch requires high-quality *de novo* sequencing data with adequate depth. In the absence of such data, this alignment-based strategy risks producing inaccurate sequence information, particularly in the pivotal CDRs. Furthermore, even advanced commercial software solutions like PEAKS AB and SuperNovo are limited to assembling a single purified monoclonal antibody sequence which significantly hinders the throughput of antibody de novo sequencing. Therefore, currently available algorithms do not satisfy the requirements for the complete and accurate assembly of full-length antibody sequences in routine applications.

In this study, we propose Fusion, an innovative *de novo* assembly method that integrates overlapping peptides and template information into complete sequences using a beam search strategy with a beam size of 20. Additionally, Fusion leverages several innovations, including a sliding window of 7 amino acids to analyze similarities and differences between template sequences, Levenshtein distance and signal consistency to guide the assembly of different antibody sequences. Our work makes the following three key contributions:

(1) Extensive experiment results demonstrate that Fusion outperforms existing antibody *de novo* assembly tools in accuracy and robustness, facilitating the acquisition of precise antibody sequence information for routine applications.
(2) Our research represents the first significant study to enhance the throughput of full-length antibody *de novo* sequence assembly sorely based on bottom-up mass spectrometry. This advancement has greatly enhanced the efficiency of monoclonal antibody research.
(3) The simultaneous analysis of multiple monoclonal antibodies under identical experimental conditions is feasible. This technique allows for the shared use of enzymes, experimental reagents, and mass spectrometry detection time, thereby reducing *de novo* sequencing costs and enhancing the economic efficiency of large-scale antibody research and development.

## 2 Results

### 2.1 Overview of Fusion

The Fusion assembler, a novel approach for antibody sequence assembly, is structured into four distinct stages (Fig. 1). Initially, it accepts unfiltered *de novo* peptide sequencing output, which includes *de novo* peptides along with sequence scores and local confidence scores for each amino acid. Stage I, known as the peptide preprocess, aims to retain a maximal number of peptide tags while eliminating peptides and amino acids with low confidence (default local confidence *<* 50). This process follows the recommendation from DeepNovo, which suggests that removing sequence contaminants by excluding peptides with a confidence score below 50 enhances the quality of the assembly [14]. The rationale behind this threshold is to guarantee both the quantity and quality of peptides used for assembly in the absence of prior knowledge. Stage II, template search, matches peptide reads to the translated germline sequences of related species contained in the IMGT to find the template for assembly, encompassing V, J and C segments. For the alignment between peptides and sequences, we utilized a mass-based alignment algorithm [19] that accounts for different combinations of amino acids potentially different lengths coinciding to the same mass (e.g., GG=N, GA=Q, etc.).

**Fig. 1.**
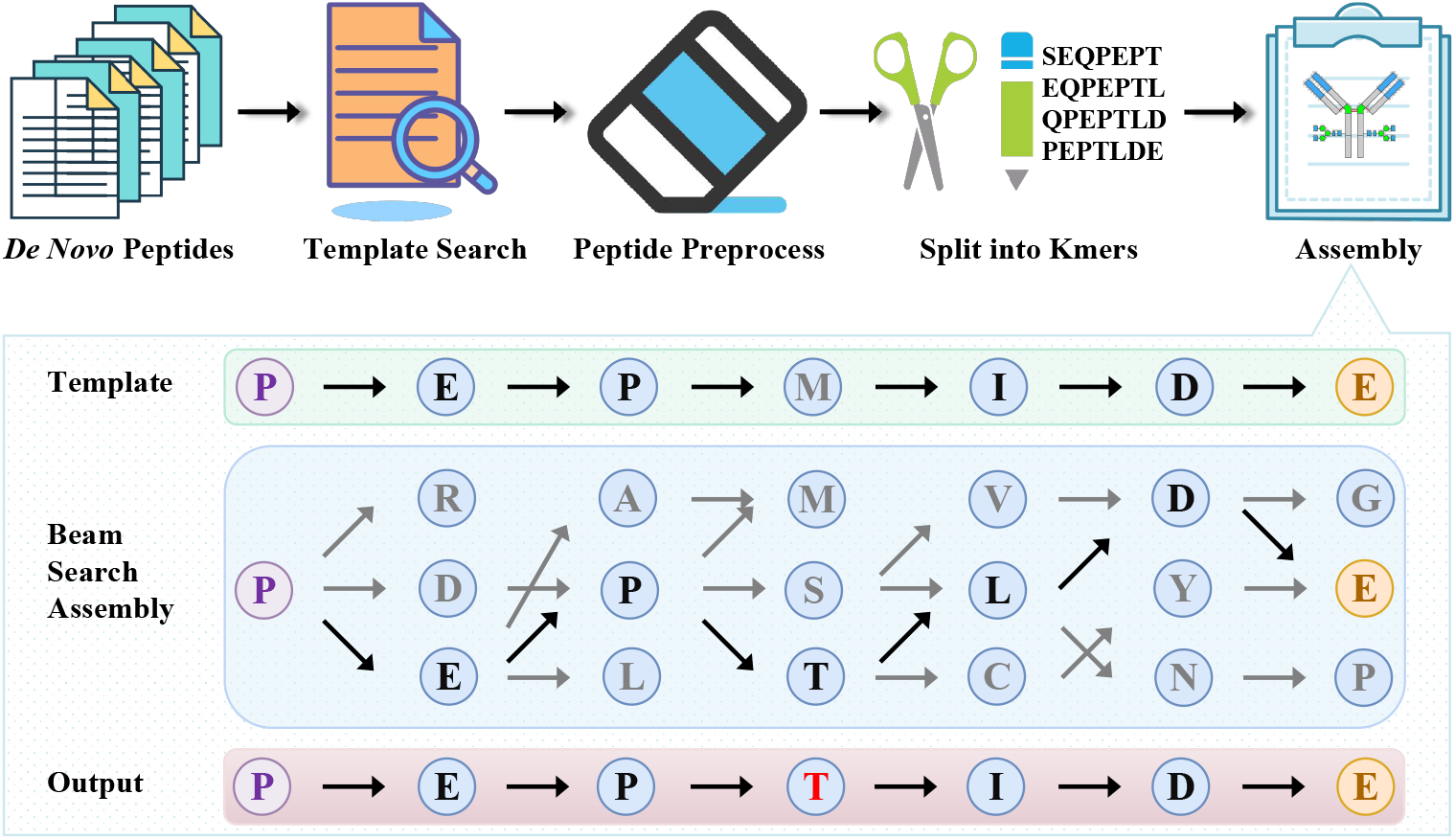
Schematic overview of Fusion.

Stage III, split into K-mers. We define a k-mer as a short peptide with k amino acids. All possible k-mers were extracted from the preprocessed peptides, and each k-mer was further split into two overlapping substrings of length *k* − 1, called left and right (*k* − 1)-mers. The left and right (*k* − 1)-mers represent nodes in the de Bruijn graph while the k-mer corresponds to a directed edge in the graph, pointing from the left to the right (*k* − 1)-mer. Based on the recommendation of Tran et al. [14], we selected *k* = 7 for the assembly of antibody sequences, a decision informed by the trade-off between the issues of sequence repetitiveness encountered with shorter k-mers and the insufficient peptide coverage associated with longer k-mers. We found that the peptides’ occurrences, positional confidence scores and the information of template provide more useful information and substantially improve the assembly quality from the de Bruijn graph. In particular, we define the confidence score of each (*k* − 1)-mer as the weighted geometric mean of its amino acids’ confidence scores. The weight of each amino acid was set to *ω* + 1 if it matched the corresponding amino acid in the template sequence; otherwise, it was assigned a default value of 1. Since a (*k* − 1)-mer can appear in multiple peptides, its weight was accumulated over the processing of all those peptides. Our formulation of the node weights is defined in the following equation:

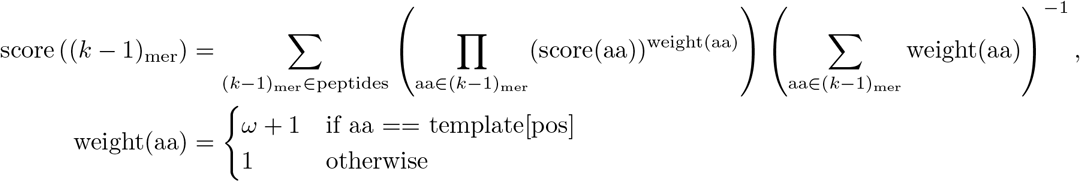

Stage IV, assembly, upon acquiring the K-mer sets and its weight scores, Fusion initiates the assembly stage by constructing the de Bruijn graph. A straightforward approach might involve choosing among a larger set of K-mer candidates generated by beam search during sequence extension. During the extension of the de Bruijn graph, we prioritize the top *N* (beam width) K-mers with the highest weight. Finally, by employing the conserved site as the target node, the sequence that ultimately achieves the highest comprehensive score is selected as the output sequence. It should be noted that template sequences are of sufficient homology to provide a correct framework for the heavy and light chains, but the exact match to the target sequence is likely not present even in the most extensive databases. This is exemplified in the lower half of Fig. 1, particularly during the extension of the 4 th amino acid, where threonine (T) is identified based on *de novo* peptide sequencing evidence. Fusion extends the contig in both forward and backward directions during the retrieval of the CDR3 region, which enables it to automatically identify and restructure D fragments for heavy chains that are composed of V-D-J-C segments, despite using only V-J-C segment information from the IMGT database.

### 2.2 *De Novo* Peptides Determination

We initially assessed the performance of the pre-trained Casanovo model using four human COVID-19 neutralizing antibody datasets (BD5514LH, and 50-200 μg S2P6LH) and three murine COVID-19 neutralizing antibody datasets (85F7, 36H6, and 2B4). Specifically, S2P6LH consists of three datasets (50 μg S2P6LH, 100 μg S2P6LH, and 200 μg S2P6LH), each corresponding to initial sample quantities of 50 μg, 100 μg, and 200 μg, respectively. From the total dataset of five proteases with HCD fragmentation, we collected peptide reads (defined as peptides with a score *≥* 50) as follows: 12,423 (BD5514LH), 15,303 (50 μg S2P6LH), 15,851 (100 μg S2P6LH), 21,967 (200 μg S2P6LH), 29,993 (85F7), 27,218 (36H6), and 32,862 (2B4). As shown in Fig. 2, the sequence coverage achieved 100% for both heavy and light chains across variable and constant domains. Moreover, the mean depth of coverage in the CDRs, variable domain, and overall chain in both heavy and light chains ranged from 56.12, 76.67, and 107.58 to 212.44, 232.25, and 283.70 (Table S1), respectively. Consequently, the *de novo* peptide reads deciphered by the pre-trained Casanovo model can be assembled into complete sequences for each antibody.

**Fig. 2.**
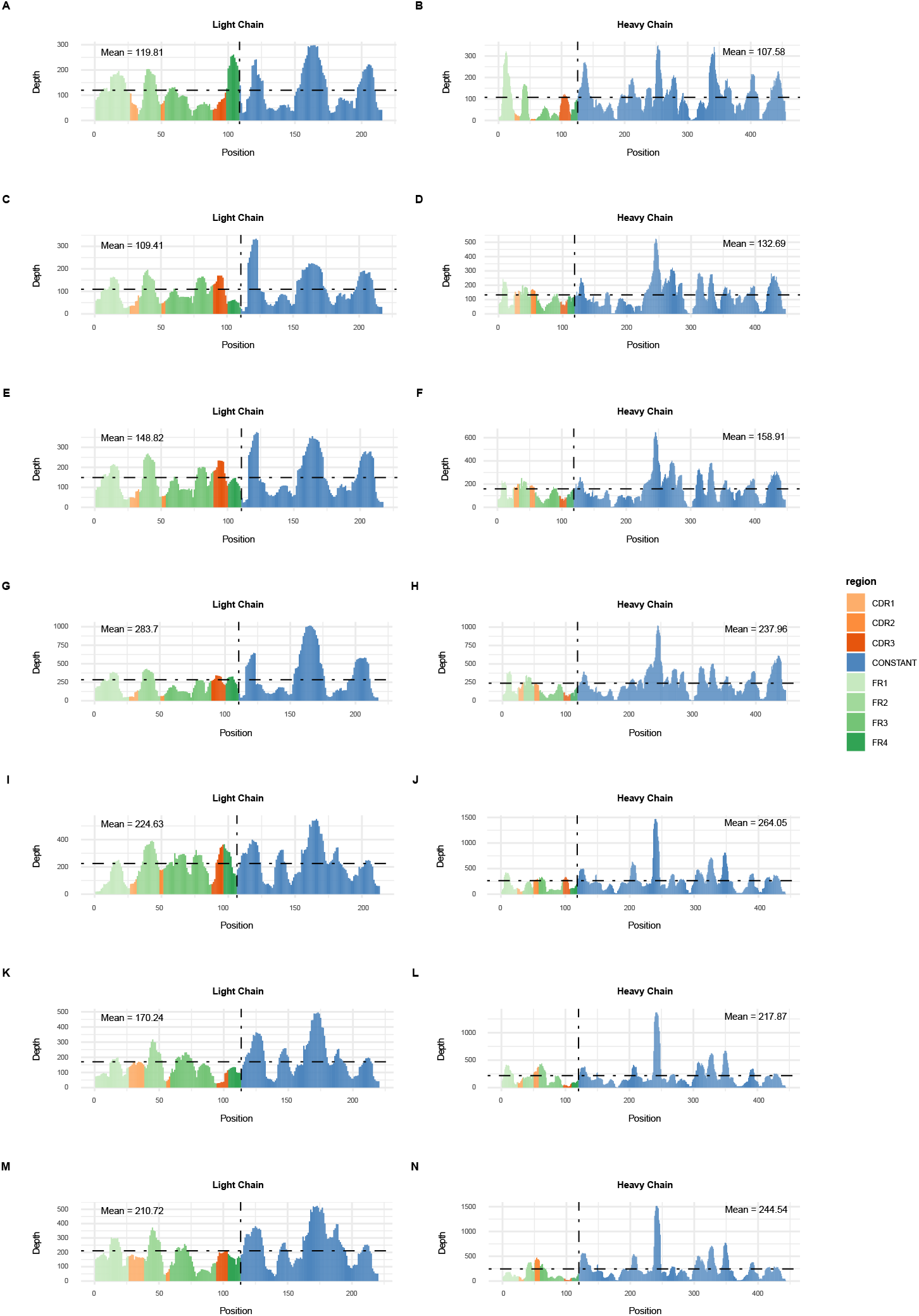
The coverage and depth of antibodies with known sequence. (A-B) BD5514LH antibody, (C-D) 50μg S2P6LH antibody, (E-F) 100μg S2P6LH antibody, (G-H) 200μg S2P6LH antibody, (I-J) 85F7 antibody, (K-L) 36H6 antibody, (M-N) 2B4 antibody.

### 2.3 Assembly Results of Known Monoclonal Antibody

Stitch is a public state-of-the-art method for antibody sequence assembly, but its accuracy varies with peptide cutoff scores, segment orders, and software versions (see Methods for details). To rigorously evaluate the assembly performance of Fusion, we compared it with Stitch’s best results. We first conducted a comparative analysis using four human antibody datasets. Stitch showed considerable variability in coverage accuracy across different antibodies. Its accurate coverage for light chains ranged from 99.54% to 100%, while for heavy chains, it ranged from 97.58% to 99.55%, as shown in Fi. 3A-D. Specifically, Stitch failed to obtain complete and accurate sequence information of the CDR3 region of the BD5514LH antibody, resulting in accurate coverage of 76.47% for the CDR region and 97.58% for the heavy chain. In contrast, Fusion exhibited more consistent and robust performance, maintaining 100% accurate coverage for both heavy and light chains, as detailed in Fig. 3A-D. This robustness is further evident when examining three S2P6LH antibody datasets across different sample quantities, where Stitch shows inconsistencies, whereas Fusion yields consistent results regardless of sample content and batch differences.

**Fig. 3.**
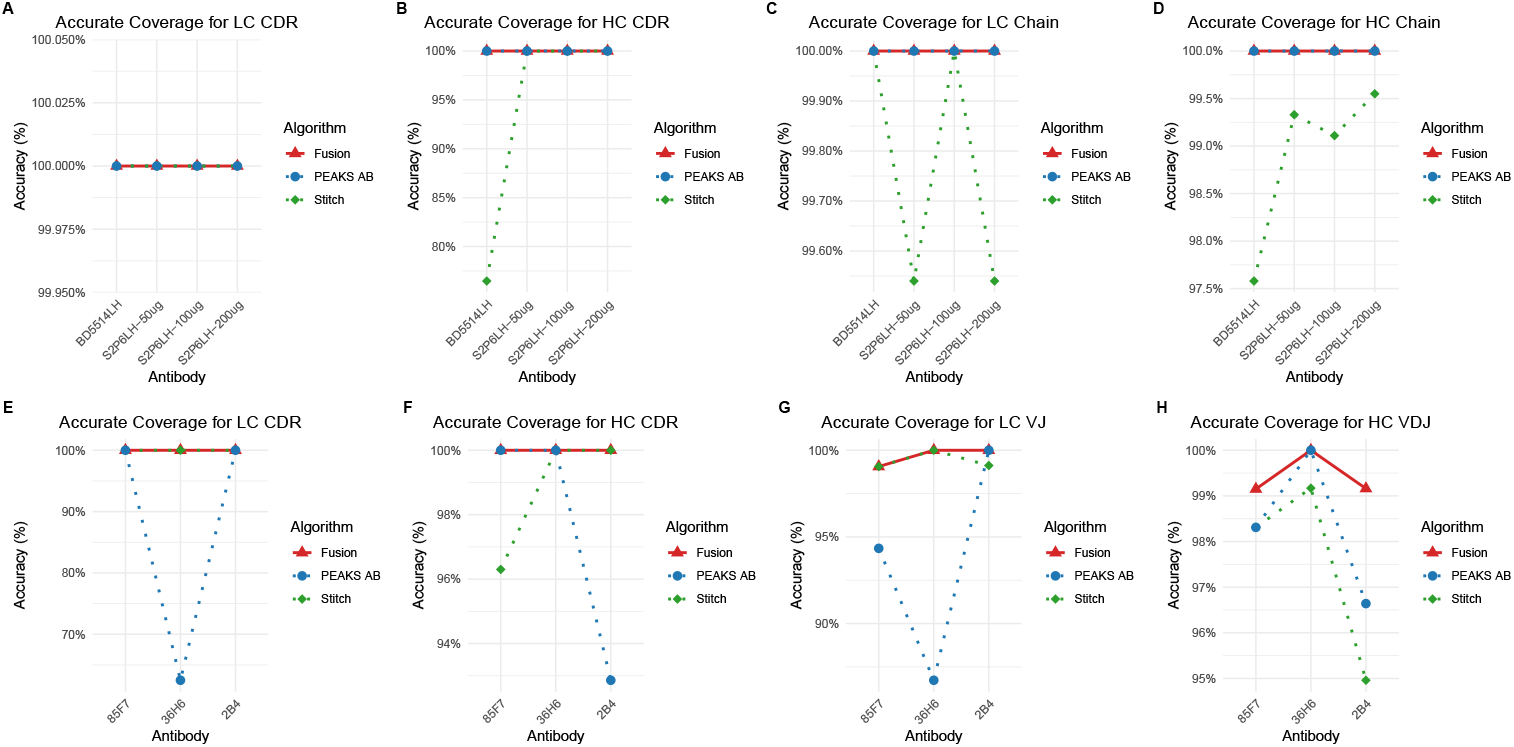
Summary of *de novo* assembly results generated by Fusion, PEAKS AB, and Stitch algorithms. (A-D) Results of human antibody datasets with known sequences. (E-H) Results of murine antibody datasets with known variable region sequences.

Next, we conducted a comparative analysis with Stitch and PEAKS AB using three murine antibody datasets with known variable region sequences. Fusion obtained the most complete and accurate sequences for both heavy and light chain variable regions, achieving 100% accurate coverage of the CDR regions for all antibodies (Fig. 3E-H). In contrast, Stitch and PEAKS AB showed considerable variability in accurate coverage across different antibodies. The accurate coverage of Stitch for light chain variable regions fluctuated between 99.06% and 100%, while for heavy chain variable regions, it ranged from 94.96% to 99.17%. More importantly, Stitch couldn’t obtain the complete heavy chain CDR3 sequence of the 85F7 antibody due to amino acid gaps. Similarly, the performance of PEAKS AB on murine antibody datasets was inconsistent, with accurate coverage for light chain variable regions fluctuating between 62.5% and 100%, and for heavy chain variable regions ranging from 92.86% to 100%. Additionally, PEAKS AB was unable to obtain the complete heavy chain CDR3 sequence of the 2B4 antibody due to amino acid errors and insertions, and it also failed to assemble the accurate light chain CDR3 sequence of the 36H6 antibody.

Last but not least, full sequence alignment of the heavy chain variable region for 85F7 and 2B4 antibodies showed differences between DNA generated sequence and mass spectrometry generated sequence. All *de novo* peptide reads supported the last amino acid in the variable region as S, not Q. This suggests that the actual accuracy of Fusion for murine antibodies is higher than the aforementioned results.

In conclusion, the comparative study highlights Fusion’s superior assembly performance over Stitch and PEAKS AB, especially in terms of accuracy and consistency across different samples and critical regions like the CDRs, making it a more reliable tool for precise antibody sequence analysis and research.

### 2.4 Application of Fusion to a commercial monoclonal antibody with undetermined sequence

An emerging therapeutic strategy in onco-immunology involves using antibodies to control tumor growth or eradicate cancer through immunotherapy. We aimed to demonstrate the utility of Fusion for therapeutic antibody research. The AFS98 monoclonal antibody effectively abrogates tumor growth by depleting macrophages and targeting the mouse colony-stimulating factor 1 receptor (CSF1R), which regulates the proliferation and differentiation of monocytic lineage cells. The AFS98 antibody was chosen as a test case and analyzed through Casanovo and Fusion to determine its sequence information. After *de novo* peptide sequencing, we collected 29,838 peptide reads from HCD spectra. Fusion successfully assembled these peptides into the complete sequences for both heavy and light chains, with mean depth of coverage in the CDRs, variable domain, and overall chain being 163.76, 156.08, and 269.77 for the heavy chain, 239.70, 231.58, and 238.33 for the light chain, respectively. Additionally, we also executed PEAKS AB software (v3.0) to analyze AFS98 antibody, which obtained longer sequences for both heavy and light chains compared to Fusion (Fig. 4A). However, the intact mass analysis indicated that the sequence assembled by Fusion more closely matches the molecular weight of AFS98 antibody, whereas PEAKS AB shows significant deviation (Table 1). Undoubtedly, Fusion is more promising in characterizing the actual sequence of AFS98 antibody. Further validation using biochemical methods is still warranted to confirm the accuracy of the sequences generated by Fusion.

**Table 1.** Summary of Intact Mass Analysis.

| Algorithm | Chain | Calculated mass (Da) | Measured mass (Da) | $\Delta$ mass (Da) |
| --- | --- | --- | --- | --- |
| PEAKS | HC | 49193.03 | 47724.89 | -1468.14 |
|  | LC | 23627.03 | 23278.19 | -348.84 |
| Fusion | HC | 47726.30 | 47724.89 | -1.41 |
|  | LC | 23288.67 | 23278.19 | -10.48 |

**Fig. 4.**
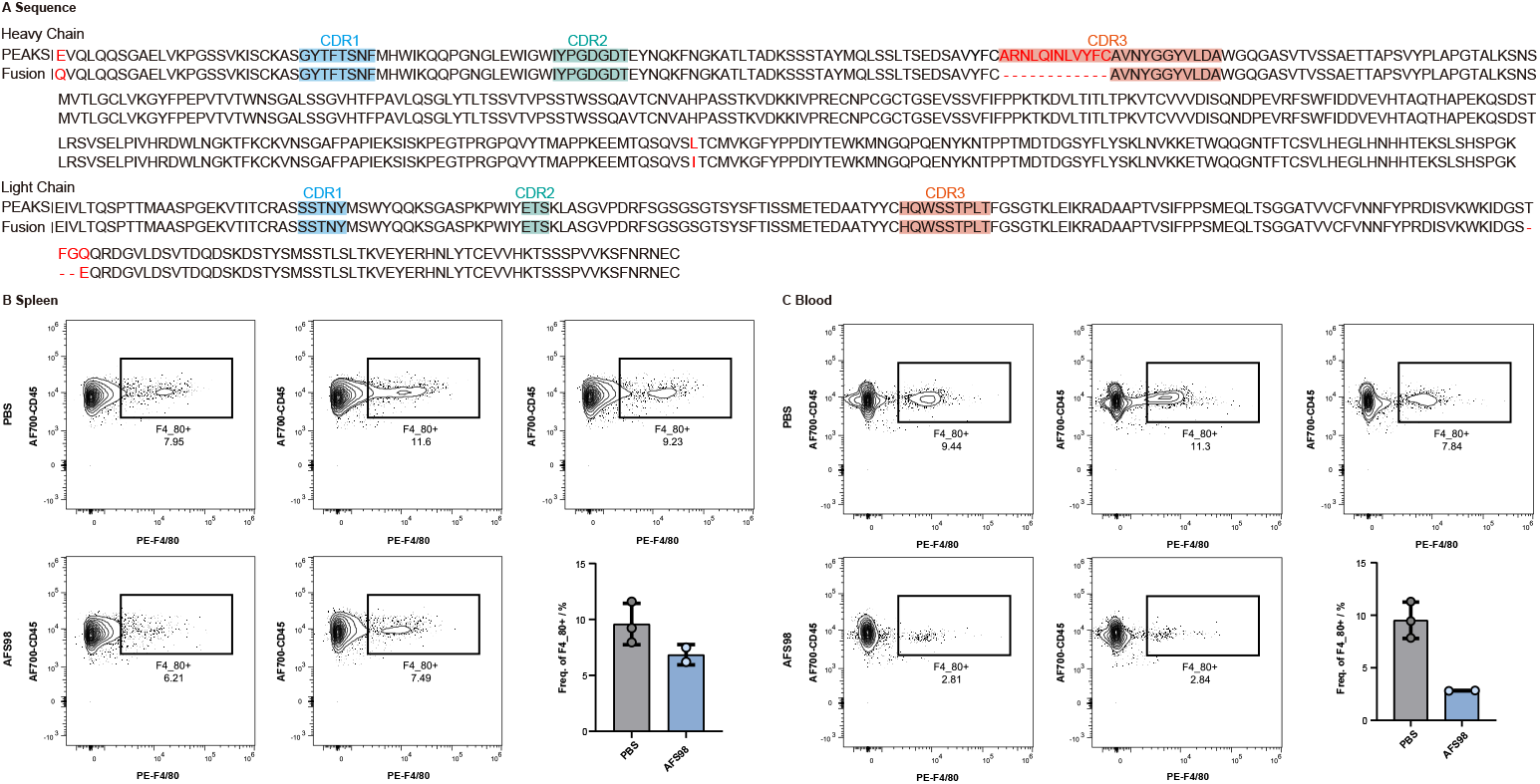
Application of Fusion to anti-mouse CSF1R antibody (AFS89). (A) The amino acid sequence of AFS89 antibody using two assembly methods, PEAKS AB and Fusion. A comparison of the efficiency in clearing macrophage in spleen (B) and blood (C) between recombinantly expressed AFS98 antibody and a blank control group using PBS buffer.

Subsequently, the sequence generated by Fusion algorithm was cloned and transfected into cells for recombinant expression, followed by affinity purification. We then administered the recombinant antibody and a commercial antibody to two groups of C57BL/6J mice, respectively. By the third day post-injection, effective clearance of macrophages in both spleen and blood was observed. Compared to control mice without macrophage depletion, the recombinant antibody achieved 28.60% and 70.35% clearance efficiency in the spleen and blood (Fig. 4B-C), respectively, demonstrating its effectiveness. This finding offers biological validation for the Fusion algorithm, underscoring its effectiveness in accurate sequence reconstruction for unknown antibodies and its potential impact on antibody research and development.

### 2.5 Assembly Results of Monoclonal Antibody Mixture

Despite the success of Fusion in assembling a single purified monoclonal antibody sequence, the feasibility of sequencing multiple monoclonal antibodies simultaneously using bottom-up mass spectrometry remains uncertain. This task presents a particular challenge due to the complexity of distinguishing and assembling sequences from a mixture of different antibodies. However, if successful, Fusion could significantly enhance the throughput of monoclonal antibody research.

To explore the feasibility of antibody mixture sequencing, we first mixed *de novo* peptide reads from three murine monoclonal antibody datasets. During the template alignment phase, we found that the top five IGHV segments contained the templates of these three antibodies and their homologous sequences, exhibiting similarities exceeding 90%. Additionally, the top three light chain V segments served as templates for these antibodies. Based on these findings, we designed a template selection procedure that sequentially adds the V segments with the highest alignment scores to the template library until the number of V segments matches the number of antibodies to be detected. During this iterative process, segments with more than 90% similarity to any already in the template library were discarded. The J segments for both heavy and light chains, which are relatively short (not exceeding 20 amino acids), were selected based on information from the V segment and peptide overlap. The C segment, being a relatively conserved sequence with few mutations, was selected based on its connection to the J segment and peptide overlap information. Thus, accurate templates for each antibody from the mixture were obtained. We then utilized a seven-amino-acid sliding window to analyze similarities and differences between the antibody template sequences. Additionally, we incorporated Levenshtein distance and signal consistency to assist in the sequence assembly of the antibody mixture. As a result, we initially achieved the assembly of the amino acid sequences for the three antibodies.

Subsequently, we applied the antibody mixture assembly protocol to real murine monoclonal antibody mixtures using bottom-up mass spectrometry as test cases. We mixed 100 μg each of the 85F7, 36H6, and 2B4 antibodies, or mixed any two of them in equal sample quantity, and performed *de novo* sequencing of these antibody mixtures separately. After completing *de novo* peptide sequencing with the pre-trained Casanovo model, we employed Fusion to assemble peptide reads from the antibody mixture into complete heavy and light chains. Remarkably, Fusion assembled the same sequences as those obtained from traditional single antibody sequencing (Fig. 5 and Fig. 6), regardless of whether the mixture contained two or three antibodies. The experiments confirmed that Fusion excelled by accurately assembling both heavy and light chains of the antibody mixture and correctly pairing them into individual monoclonal antibodies. Such accomplishment reflects a level of assembly performance comparable to that observed in traditional single monoclonal antibody assembly, highlighting Fusion’s exceptional accuracy and flexibility.

**Fig. 5.**
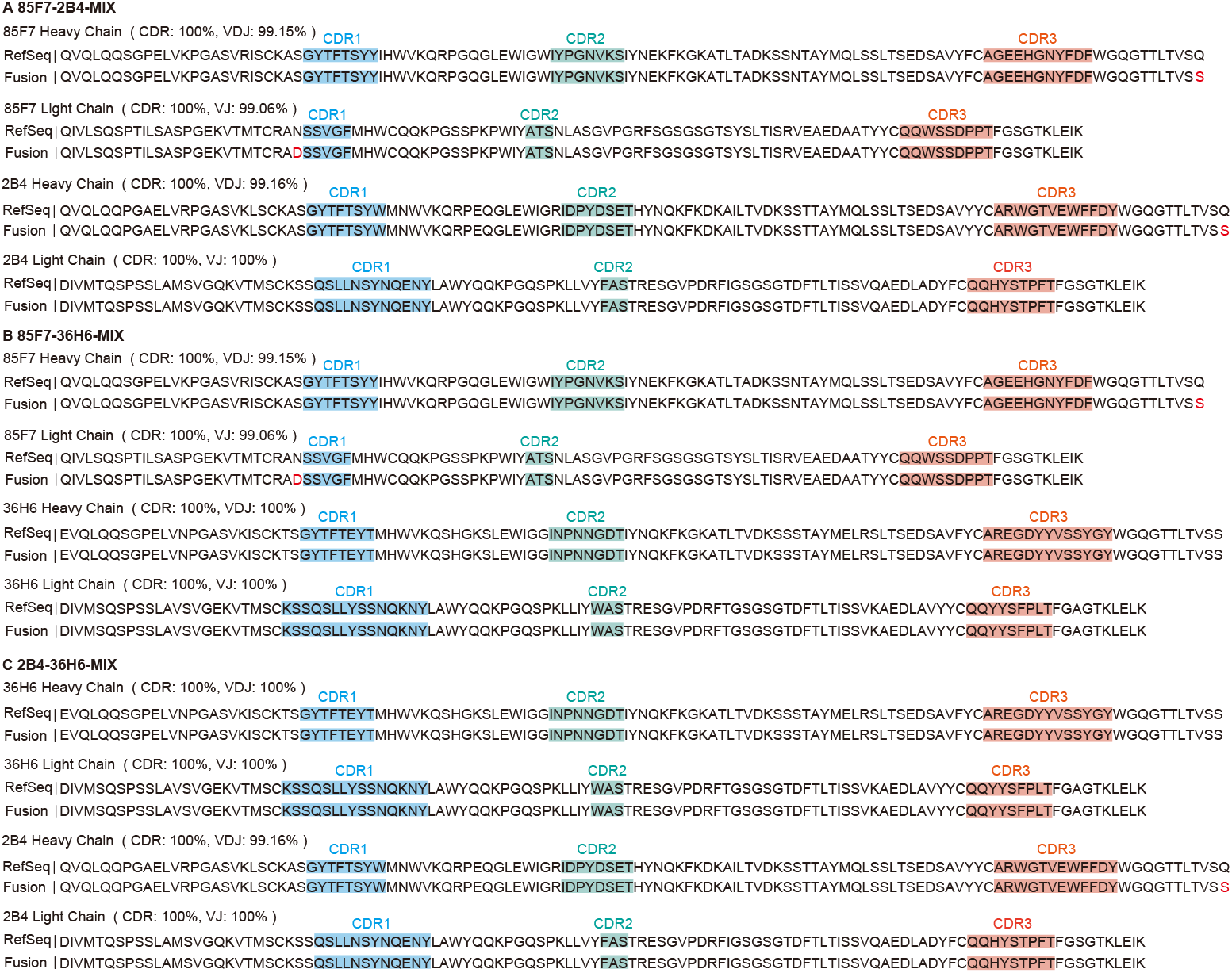
Assembly results for antibody mixture containing two monoclonal antibodies. (A) 85F7-2B4-MIX, (B) 85F7-36H6-MIX, (C) 2B4-36H6-MIX.

**Fig. 6.**
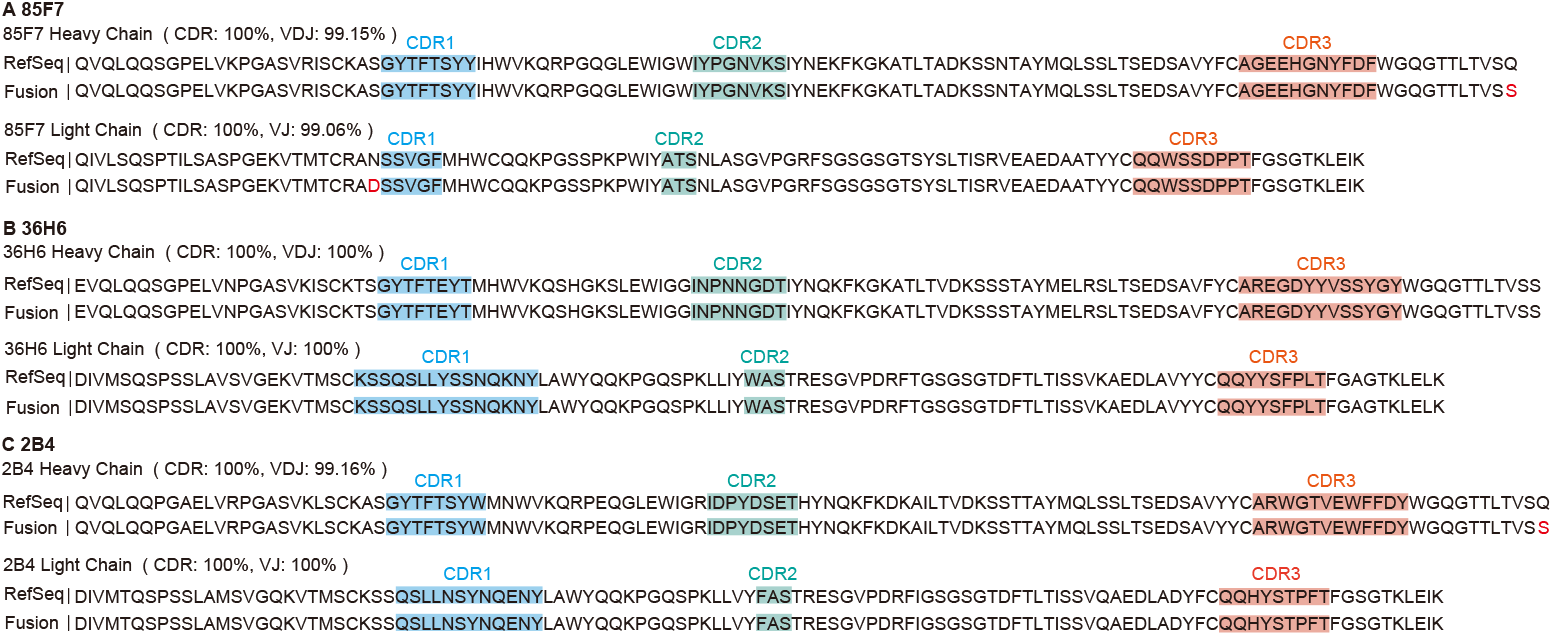
Assembly results for antibody mixture containing three monoclonal antibodies.

This methodology significantly enhances throughput and cost-effectiveness. By employing Fusion to simultaneously assemble multiple monoclonal antibodies within a single experimental framework, the need for separate, sequential processes is eliminated. Such optimization conserves time, reduces reagent consumption, and lowers equipment-related operational costs. Consequently, Fusion represents a major advancement in antibody research methodologies, enhancing efficiency and reducing expenses.

## 3 Discussion

A contemporary application of antibody *de novo* sequencing based on bottom-up mass spectrometry involves determining the sequence and its variants of an innovator biologic to develop a biosimilar. With $67 billion worth of biopharmaceutical product patents expired by 2020, the market for biosimilar production is growing, and *de novo* sequencing is the first step in determining and verifying the innovator amino acid sequence. However, the throughput remained a challenge, particularly in the biopharmaceutical industry where time constraints are a major concern. Previously, we developed a sample preparation method, referred to as SP-MEGD (Single-Pot and Multi-Enzymatic Gradient Digestion), which engenders numerous missed cleavage events and produces various peptide products of diverse lengths with overlapping stretches of residues, enabling efficient database-free de novo sequencing. Additionally, a variety of deep learning tools are capable of de novo sequencing individual spectra and providing adequate peptide-level information. Consequently, the key challenge, and thus the bottle neck remains within the field of bioinformatics, particularly in the task of sequence assembly.

With our efforts, Fusion algorithm is capable of assembling peptide sequences from mixtures containing two or three monoclonal antibodies. Fusion integrates several algorithmic innovations, providing hands-free operation and delivering robust results in high-throughput fashion. This capability significantly reduces the necessity for conducting separate de novo sequencing experiments for each monoclonal antibody, thereby diminishing the duration, expense, and reagent consumption of mass spectrometry detection. As a result, the efficiency of monoclonal antibody research is markedly improved. Moving forward, we aim to further refine the Fusion algorithm, enhancing its utility in facilitating more extensive antibody research and development initiatives.

Besides, Fusion can be used to assembly antibody sequence in traditional single monoclonal antibody sequencing fashion. To rigorously evaluate the assembly performance of Fusion, we constructed large-scale comprehensive antibody datasets with known sequences. Fusion achieves 100% accuracy for multiple known human antibodies and over 99% accuracy for murine antibodies with known variable region sequences, representing the highest accuracy in its class. Moreover, we have effectively demonstrated the utility of Fusion for therapeutic antibody research by conducting a biological validation of the recombinantly expressed AFS98 antibody.

For a given species, Fusion can automatically assemble antibody sequences by simply inputting *de novo* peptides along with sequence scores and local confidence scores for each amino acid. Therefore, Fusion is well-suited for characterizing any unknown antibody of known model species. However, achieving complete de novo assembly of novel or unknown antibody sequences without templates remains a significant challenge due to limitations in peptide fragmentation, coverage, and spectral interpretation ambiguities. To date, no assembly algorithm has really achieved this. Nonetheless, with ongoing advancements, the Fusion algorithm may, in the future, enable the complete assembly of antibody sequences in a template-independent manner, even for non-model organisms. Additionally, in theory, Fusion can be adapted to any *de novo* peptide sequencing software. For example, Fusion also obtained complete antibody sequence information based on *de novo* peptide reads from PEAKS AB.

## 4 Methods

### 4.1 Monoclonal Antibodies and Monoclonal Antibody Mixture Sample Preparation

An anti-mouse CSF1R monoclonal antibody AFS98 was purchased from BioXCell (BE0213). Three human COVID-19 neutralizing antibodies (S2P6LH, BD5514LH, and BD5840LH) and three murine COVID-19 neutralizing antibodies (85F7, 36H6, and 2B4) were expressed by the ExpiCHO Expression System (Thermo Fisher Cat#A29133) and subsequently purified via a ProteinA column (Cytiva). The sequence information of COVID-19 neutralizing antibodies is listed in Table 1. Specifically, three murine antibodies with amino acid sequences known only for their variable regions rather than complete chains.

Following purification, the Fc-tagged proteins were pooled and concentrated using a Protein Quantification Kit (Thermo Fisher Cat#23225). The overall sample preparation mainly includes three steps. Firstly, 100 μg sample was denatured and reduced at 60°C for 30 min with 6 M guanidine hydrochloride (Gu.HCl) and 20 mM dithiothreitol (DTT), followed by alkylation (40 mM iodoacetamide, IAA) in the dark for 30 min at room temperature. For protein digestion, five enzymes (trypsin, chymotrypsin, pepsin, elastase, and ASPN) were respectively added to the sample in a ratio of 1:20 (w/w). Proteins were digested by an incubation at 900 rpm for 6 h at 37°C. After digestion, the reaction mixture was quenched by trifluoroacetic acid (TFA, 10% v/v) for 30 min at 37°C. The supernatant was desalted using Sep-Pak C18 Vac cartridges (Waters), lyophilized to dryness, and stored at -20°C or resuspended in 0.1% formic acid (FA) for nano-LC-MS/MS analysis.

### 4.2 LC-MS/MS

The digested peptides were separated by online reversed-phase chromatography on a Vanquish Neo UHPLC coupled to a Thermo Scientific Orbitrap Eclipse mass spectrometer. The peptides were separated on a 75 μm *×* 50 cm long column (2 μm id) at a flow rate of 300 nL/min for 70 min. Full MS scans were acquired in an Orbitrap detector in the range of *m/z* 350-2000, at a resolution of 60000 (at *m/z* 200), with an automatic gain control (AGC) of 4e5 and a maximum injection time of 50 ms. Data-dependent MS2 acquisition was also performed in the Orbitrap, using a topspeed cycle time of 1.5s at a 30000 resolution (at *m/z* 200). The precursors were fragmented by stepped high-energy collision dissociation (HCD) as well as electron-transfer high-energy collision dissociation (EThcD). The HCD fragmentation employed three stepped normalized collision energy (NCE) settings of 27%, 35%, and 40%, with a maximum injection time of 250 ms. For EThcD fragmentation, the activation type was switched to charge-dependent ETD followed by HCD fragmentation with a supplemental activation-NCE of 30%. Finally, 11 bottom-up mass spectrometry datasets were generated for this study, based on traditional single monoclonal antibody sequencing (Table 2) or monoclonal antibody mixture sequencing (Table 3).

**Table 2.** The informations of monoclonal antibody datasets.

| Antibody Dataset | LC / VJ (AA) | HC / VDJ (AA) |
| --- | --- | --- |
| BD5514LH | 215 | 455 |
| 50 $\mu\text{g}$ S2P6LH | 217 | 447 |
| 100 $\mu\text{g}$ S2P6LH | 217 | 447 |
| 200 $\mu\text{g}$ S2P6LH | 217 | 447 |
| 85F7 | 106 | 118 |
| 36H6 | 113 | 120 |
| 2B4 | 113 | 119 |

Notes: Except for S2P6LH, which had three different sample quantities, the sample quantity for each other antibodies was 100 $\mu\text{g}$ .

**Table 3.** The informations of monoclonal antibody mixture datasets.

| Mixture | Composition |  |  |
| --- | --- | --- | --- |
| 2-Mix | 85F7-2B4-MIX | 85F7-36H6-MIX | 2B4-36H6-MIX |
| 3-Mix | 85F7-2B4-36H6-MIX |  |  |

Notes: Mixed two or three monoclonal antibody in equal sample quantity, each 100 $\mu\text{g}$ .

### 4.3 Deep Learning Model for *De Novo* Sequencing of MS/MS Spectra

Casanovo [15] is the model using the Transformer architecture for *de novo* sequencing, and it preprocessed MS2 using sine and cosine waves. We trained the Casanovo model (v3.2.0) for 20 epochs with its pre-defined parameters on the MassIVE KB v1 dataset [20], which encompasses over 2.1 million precursors from 19,610 proteins. This dataset was meticulously assembled from more than 31 TB of human data, sourced from 227 public proteomics datasets. Notably, it maintains rigorous false discovery rate controls, ensuring a comprehensive and robust training environment for our model. We split the spectra into training and validation sets at a ratio of 98:2 while making sure that the split data sets did not share any common peptides. During training, 10 ppm for the precursor mass tolerance and 0.02 Da for the fragment mass tolerance were specified. Carbamidomethylation of cysteine (C + 57.02 Da) was set as a fixed modification, while oxidation of methionine (M + 15.99 Da), deamidation of asparagine, glutamine (N + 0.98 Da and Q + 0.98 Da) and Pyroglutamate formation from glutamine (Q-17.03 Da) were set as variable modifications.

Each instrument vendor employs its own file formats to store results from MS/MS experiments, requiring conversion to open-format files for compatibility with the pretrained Casanovo model. We used ProteoWizard software [21] to reformat the raw MS/MS data files of each antibody dataset to Mascot Generic Format (MGF). MGF files store the *m/z* and intensity pairs of multiple mass spectra in a single text format. Afterward, Casanovo deciphers amino acid sequences by interpreting the mass variations observed in the MS/MS peaks.

### 4.4 Assembly of Stitch for Known Monoclonal Antibody

Stitch has recently emerged as a state-of-the-art public method for antibody sequence assembly. Users are required to define two main parameters: the cutoff score for peptide reads and the recombined segment orders. The typical recombined segment orders are IGHV*IGHJ IGHC for the heavy chain and IGLV*IGLJ IGLC for the light chain. The asterisk (*) represents a gap between adjacent segments, which is extended by overhanging peptide reads. We have chosen cutoff scores of 95, 90, 85, and 50 for peptide reads. Scores of 95, 90, and 85 are based on the default parameters from Stitch’s paper or batch file examples, while 50 is used as the threshold in Fusion. We conducted tests using both the stable version Stitch1.4.0 and the latest version Stitch1.5.0 with these peptide read cutoff scores, common recombined segment orders, and other default parameters.

During the first run (Fig. 7A-D), all assembly results for the light chain contained gaps (Fig. 7A, C), because the actual length of light chain CDR3 for BD5514LH and S2P6LH antibodies exceeds the template. Therefore, we modified the recombined segment order of the light chain to IGLV*IGLJ IGLC for these two antibodies. When we refined the Stitch run to force selection of IGLV*IGLJ IGLC for the light chain recombined segment order, almost all the CDRs’ accuracy of human antibody datasets was improved to 100%, except for the BD5514LH antibody run by Stitch1.4.0 with a 95 peptide cutoff score (Fig. 7E). Consequently, the overall accuracy of the light chain also improved (Fig. 7G). In the results of four human antibody datasets, different peptide cutoff score settings significantly impacted the outcomes, both in terms of CDR accuracy and the accuracy of the entire chain (Fig. 7E-H). This phenomenon was also observed in three murine antibody datasets (Fig. 7I-L). Notably, the actual length of the 85F7 antibody CDR3 is shorter than its template, yet all results assembled a CDR3 sequence of the same length as the template (Table S2). Moreover, sometimes Stitch1.4.0 achieved better assembly results, while sometimes Stitch1.5.0 did.

**Fig. 7.**
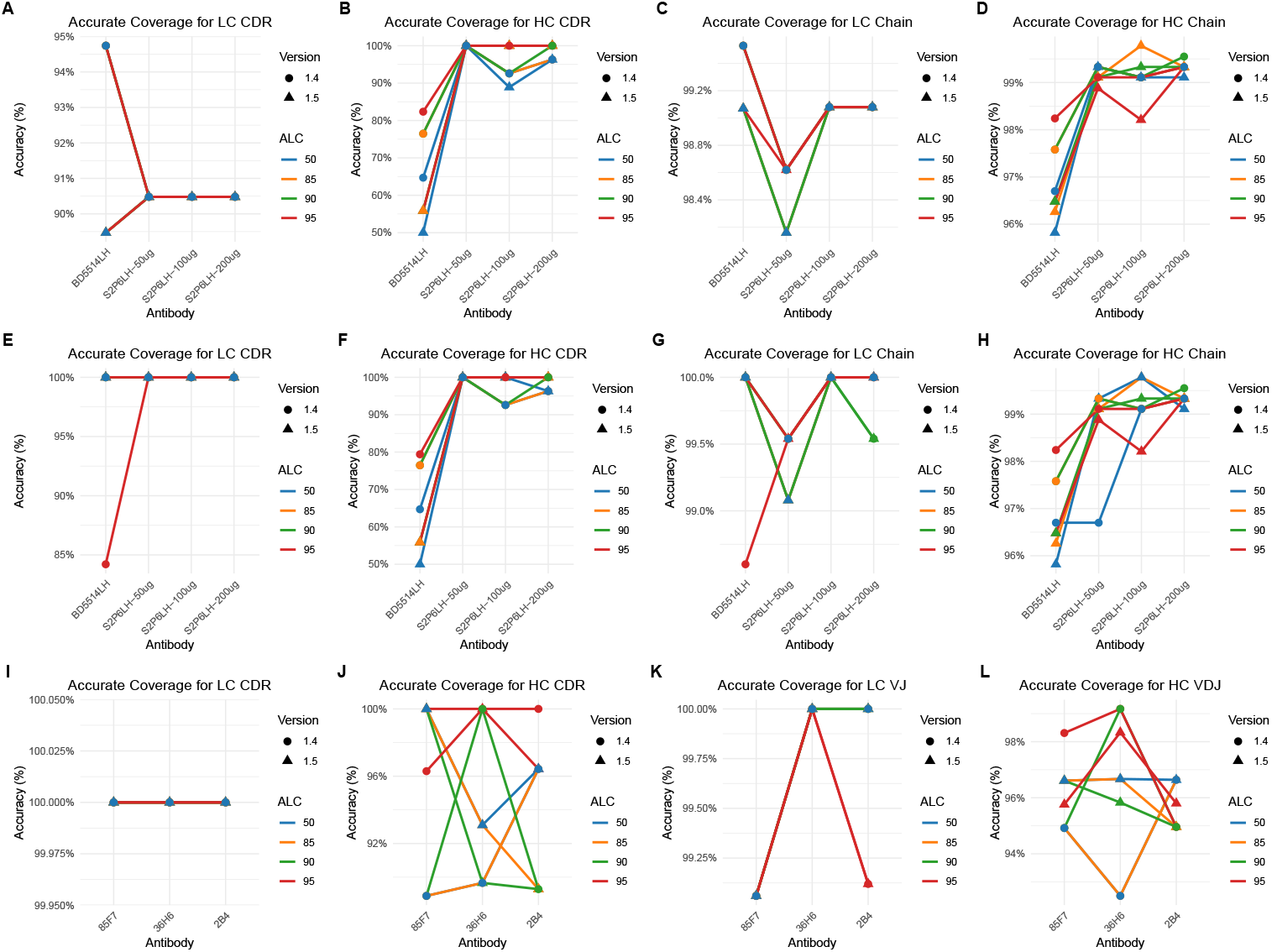
The assembly results of Stitch for known monoclonal antibody. (A-D) The result of first run for human antibodies with known sequence. (E-H) The modified result of first run for human antibodies with known sequence. (I-L) The result of murine antibodies with known variable region sequence.

In summary, different peptide cutoff scores and recombined segment orders impact the accuracy of assembly for Stitch. Different software versions also influence the assembly results. Without expert knowledge, these factors can be detrimental to the users. To rigorously evaluate the assembly performance of the Fusion algorithm, we compared its results with the best results from Stitch.

### 4.5 Germline sequences

The integration of homologous sequences significantly enhances sequence assembly accuracy by mitigating error signal effects. Germline sequences for the V, J, and C segments, sourced from the IMGT database [22] (https://www.imgt.org/IMGTrepertoire/Proteins/), undergo filtering to remove amino acid level duplicates, resulting in a catalogue of unique, nonredundant sequences. The simulation of V-(D)-J-C recombination and the subsequent full permutation of V, J, and C fragments further refine this process. Specifically, genes for the heavy chain reside within the immunoglobulin heavy-chain locus (IGH), and those for the light chain within the immunoglobulin kappa (IGK) or lambda (IGL) loci. For light chains, combinations are constrained to sequences of the same type, producing exclusively IGKV-IGKJ-IGKC or IGLV-IGLJ-IGLC sequences, thereby ensuring a methodological approach that reflects the genetic intricacies of antibody diversity and facilitates the generation of biologically pertinent recombinant sequences. Templates for the D-segment are not retrieved from IMGT due to their typical brevity and variability, but they can be automatically reconstructed using our Fusion algorithm.

### 4.6 Evaluation metrics

Sequencing coverage was calculated as the percentage of amino acids of the antibody sequence that were covered by assembled contig. Sequencing accuracy was calculated as the percentage of all annotated sequence calls that were labeled correct. To evaluate the performance of sequence assembly comprehensively, we employed the metric of accurate coverage, defined as the percentage of amino acids of the antibody sequence that were correct covered by assembled contig, equal to the product of coverage and accuracy. For the alignment between assembled sequences and actual antibody sequences, we utilized the global Needleman-Wunsch algorithm [23] in conjunction with the BLOSUM60 matrix to achieve the most intuitive results. All alignment results are presented in the Supplementary Material.

### 4.7 Discriminating Isoleucine from Leucine

The EThcD-based methodology facilitates the discrimination between isomeric leucine and isoleucine residues by leveraging specific radical losses. Specifically, a leucine-associated z-ion releases an isopropyl radical, leading to a mass reduction of 43.0548 Da and the subsequent formation of the corresponding w-ion. Conversely, isoleucine experiences the loss of an ethyl radical from its z-ion, resulting in a mass decrease of 29.0391 Da and the generation of the pertinent w-ion. We performed a database search on EThcD spectra using OMSSA [24], utilizing the assembled sequences as our protein database. OMSSA database search was executed at a precursor tolerance of 10 ppm and fragment mass tolerance of 0.02 Da. Carbamidomethylation of cysteine (C + 57.02 Da) was set as a fixed modification, while oxidation of methionine (M + 15.99 Da), deamidation of asparagine and glutamine (N + 0.98 Da and Q + 0.98 Da) were set as variable modifications.

Furthermore, the differentiation process of leucine and isoleucine residues can be enhanced by considering the cleavage specificity of enzymes such as chymotrypsin and pepsin, which selectively cleave at leucine residues, not isoleucine, thereby offering an additional layer of specificity in identification.

In its application, the Fusion algorithm identifies isoleucine or leucine residues by integrating template sequence information with experimental evidence and the detection of diagnostic w-ions, thereby enhancing the precision of isomer discrimination.

## Author contributions

W.J., Y.X., and J.X. contributed equally to this work. This manuscript was written through the contributions of all authors. All authors have approved the final version of the manuscript.

## Notes

The authors declare no conflict of interest.

## Acknowledgements

This work was supported by the Scientific Research Foundation of State Key Laboratory of Vaccines for Infectious Diseases, Xiang An Biomedicine Laboratory (2023XAKJ0100081).

